# Separating overlapping bat calls with a bi-directional long short-term memory network

**DOI:** 10.1101/2019.12.15.876656

**Authors:** Kangkang Zhang, Tong Liu, Shengjing Song, Xin Zhao, Shijun Sun, Walter Metzner, Jiang Feng, Ying Liu

**Affiliations:** Jilin Provincial Key Laboratory of Animal Resource Conservation and Utilization, Northeast Normal University, Changchun, China; School of Environment, Northeast Normal University, Changchun, China; Department of Integrative Biology and Physiology, University of California, Los Angeles, California, USA; Collage of Animal Science and Technology, Jilin Agricultural University, Changchun, China

## Abstract

Acquiring clear and usable audio recordings is critical for acoustic analysis of animal vocalizations. Bioacoustics studies commonly face the problem of overlapping signals, but the issue is often ignored, as there is currently no satisfactory solution. This study presents a bi-directional long short-term memory (BLSTM) network to separate overlapping bat calls and reconstruct waveform audio sounds. The separation quality was evaluated using seven temporal-spectrum parameters. The applicability of this method for bat calls was assessed using six different species. In addition, clustering analysis was conducted with separated echolocation calls from each population. Results showed that all syllables in the overlapping calls were separated with high robustness across species. A comparison between the seven temporal-spectrum parameters showed no significant difference and negligible deviation between the extracted and original calls, indicating high separation quality. Clustering analysis of the separated echolocation calls also produced an accuracy of 93.8%, suggesting the reconstructed waveform sounds could be reliably used. These results suggest the proposed technique is a convenient and automated approach for separating overlapping calls using a BLSTM network. This powerful deep neural network approach has the potential to solve complex problems in bioacoustics.

**Author summary:** In recent years, the development of recording techniques and devices in animal acoustic experiment and population monitoring has led to a sharp increase in the volume of sound data. However, the collected sound would be overlapped because of the existence of multiple individuals, which laid restrictions on taking full advantage of experiment data. Besides, more convenient and automatic methods are needed to cope with the large datasets in animal acoustics. The echolocation calls and communication calls of bats are variable and often overlapped with each other both in the recordings from field and laboratory, which provides an excellent template for research on animal sound separation. Here, we firstly solved the problem of overlapping calls in bats successfully based on deep neural network. We built a network to separate the overlapping calls of six bat species. All the syllables in overlapping calls were separated and we found no significant difference between the separated syllables with non-overlapping syllables. We also demonstrated an instance of applying our method on species classification. Our study provides a useful and efficient model for sound data processing in acoustic research and the proposed method has the potential to be generalized to other animal species.

## Introduction

The structural identification of vocal units is essential in animal acoustic studies for sound feature analysis, sound emitter recognition, and species identification and monitoring. However, wild animal monitoring, both in the field and in the laboratory, often involves problems caused by the overlapping of different vocal units in time and frequency space, which prevents the components from being suitable for parameter analysis. As a result, the separation of overlapping sounds is an important task in bioacoustic signal processing. However, existing analysis software often struggles to process overlapping calls and previous research on the acoustic identification of animals primarily focuses on extracting target signals from background noise for species classification or population monitoring [1-4]. The process of separating overlapping calls from mixed sounds has received little attention to date and researchers conventionally abandon sounds that overlap in both time and frequency, requiring an extension of the experimental period to obtain sufficient non-overlapping recordings [5, 6]. As such, an effective method for successfully and automatically separating overlapping calls would be of significant interest and benefit to animal researchers.

Previous studies using deep neural networks have produced promising results for automated sound recognition in complex acoustic environments for animal species recognition and classification [6-8]. However, in this study, we consider the more difficult task of separating different types of syllables from overlapping calls and reconstructing sound waves from these separated signals. Existing techniques used for animal sound separation often require prohibitive quantities of labelled data. For example, multiple-instance machine learning (MIML) algorithms were proposed for use in sound feature extraction and species identification in birds [1]. However, this technique requires a cropped mask of a signal segment (without overlap) in order to extract each syllable.

Deep learning networks have been applied to bioacoustic studies but have primarily been used for classification. For instance, convolutional bidirectional recurrent neural networks (CBRNNs) have been used to identify the presence of bird calls in audio samples [4]. Acoustic features were learned by the network (a classifier) and the presence or absence of a bird call was output as an indicator. Convolutional neural networks (CNNs) have been used to predict the presence of a search-phase bat echolocation call in spectrograms. This binary classification problem was used to detect the presence of bats [2]. To our knowledge, the use of deep learning techniques to separate animal calls that overlap in both time and frequency space has yet to be reported.

Multiple studies have been conducted using deep learning-based supervised speech separation with humans. Early systems included shallow models that performed a linear transformation of given mixture features during the prediction time interval. This has included Gaussian mixture models [9], support vector machines [10], and non-negative matrix factorization [11]. However, in real-world scenarios, the mapping relationship between mixture signals and sources is typically a nonlinear transformation. Nonlinear models, such as deep neural networks (DNNs), are therefore highly applicable because of their ability to identify nonlinear structures in audio signals [12-14]. Additionally, recurrent neural networks (RNNs) that exhibit the temporal behavior of a time sequence can be trained to predict time-frequency masks for target signals and separate sources from a mixed waveform [15]. Specifically, long short-term memory (LSTM) networks, a variation of RNN models that exhibit strong learning capabilities and simple construction, have been widely used for word and continuous speech recognition [16-18]. By concatenating two separate LSTM networks, bidirectional LSTMs (BLSTMs) can predict each element of a sequence based on past and future context and can naturally account for the temporal dynamics of speech. These models are typically faster and more accurate than standard RNNs in frame-by-frame phoneme classification [19]. In addition, the BLSTM network can compensate for exploding and vanishing gradient issues that can occur during the training of standard RNN models [20]. At present, BLSTMs have achieved state-of-the-art performance for speech recognition [14, 21], natural language processing [22, 23], and speaker-independent speech separation [24]. As such, a BLSTM model was selected in this study for overlapping bat call separation.

Echolocating bats have two vocal repertoires, stereotypical echolocation calls for orientation and a variety of communication calls for social activities [25-27]. Recordings from both field and laboratory studies indicate that utterances from individual bats often overlap in both time and frequency, which provides an excellent template for research on overlapping sound separation in animals. The primary objective of this study is to develop a technique for separating two target signals (echolocation and socialization calls) from mixtures of acoustic sounds. Although deep leaning has been employed in the acoustic classification of multiple species, including nonhuman primates [28], birds [4], whales [5], and bats [2, 3], the goal of the present study is distinct from these previous cases in which deep neural networks were primarily used as classifiers.

Both overlapping and non-overlapping calls (of both echolocation and communication types) were recorded from each of the collected bat species studied in our previous work. We developed a BLSTM network and used the recorded non-overlapping calls to train the model. Recorded overlapping calls were input to the trained model and separated. Independent sound files were then reconstructed for each separated signal. The correctness of these separated signals was measured by comparing the temporal-spectrum parameters between separated calls and the initially recorded (non-overlapping) calls from each species. Finally, clustering analysis was conducted to classify the bats using separated echolocation calls, which provided a practical application of the proposed technique.

## Results

The proposed algorithm performed well and achieved high accuracy in separating overlapping calls for each of the six species. The BLSTM model was iteratively trained until the training and validation losses reached a minimum. Loss is a summation of errors made with each sample in the training or validation sets and measures how well the model adapts during optimization. Training loss for this model decreased significantly in the first epoch. The validation loss function tended toward an asymptotic value, indicating the training algorithm had converged (S2 Fig). The BLSTM model converged slightly faster when training with CF bat samples (as opposed to FM samples).

All echolocation and communication calls in the overlapping signals were correctly extracted during the separation procedure, regardless of their pulse duration or energy characteristics (see Table 1 and Fig 1). In addition, low-intensity FM components in echolocation pulses were successfully extracted from three CF bat species (Figs 1d, 1e, and 1f).

**Table 1.**
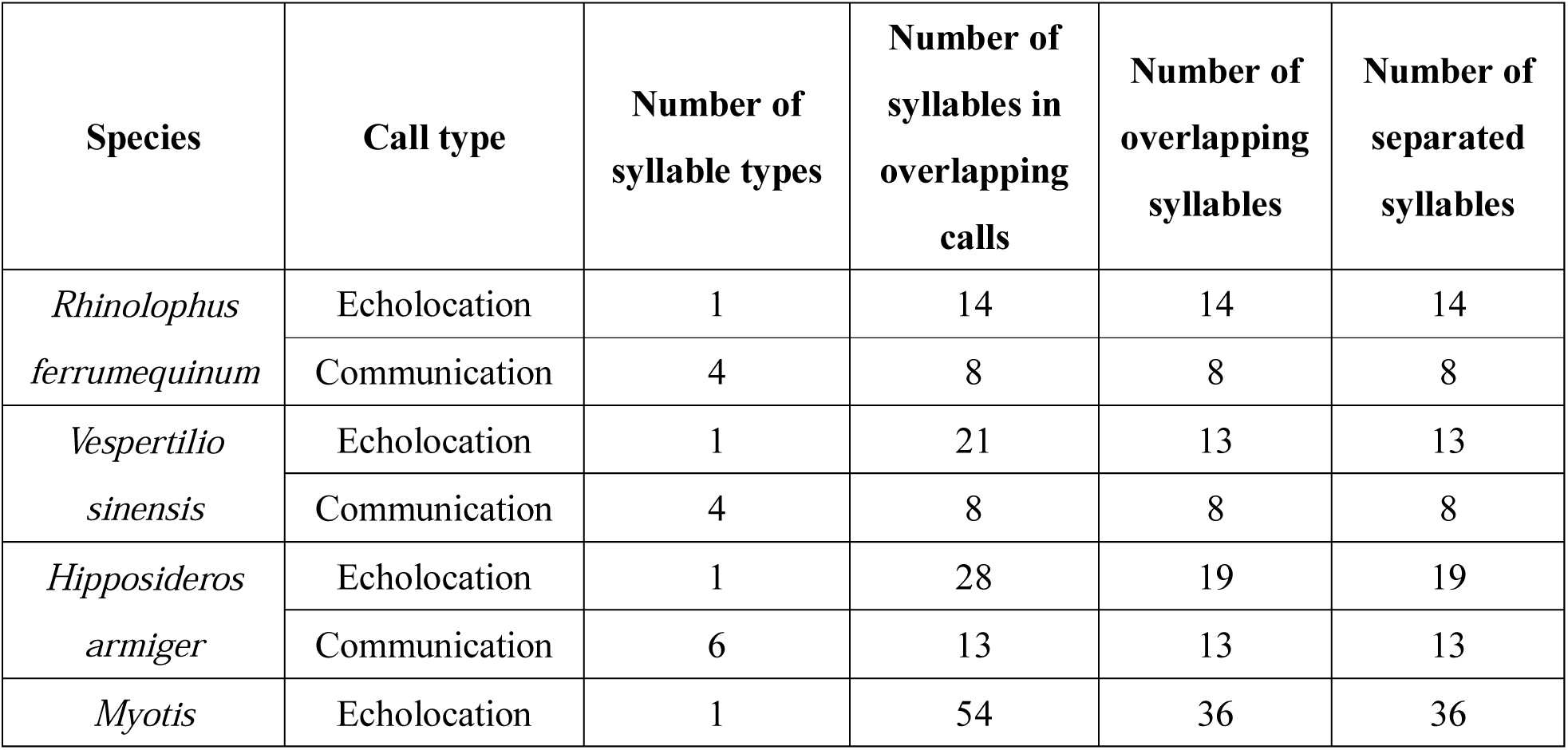
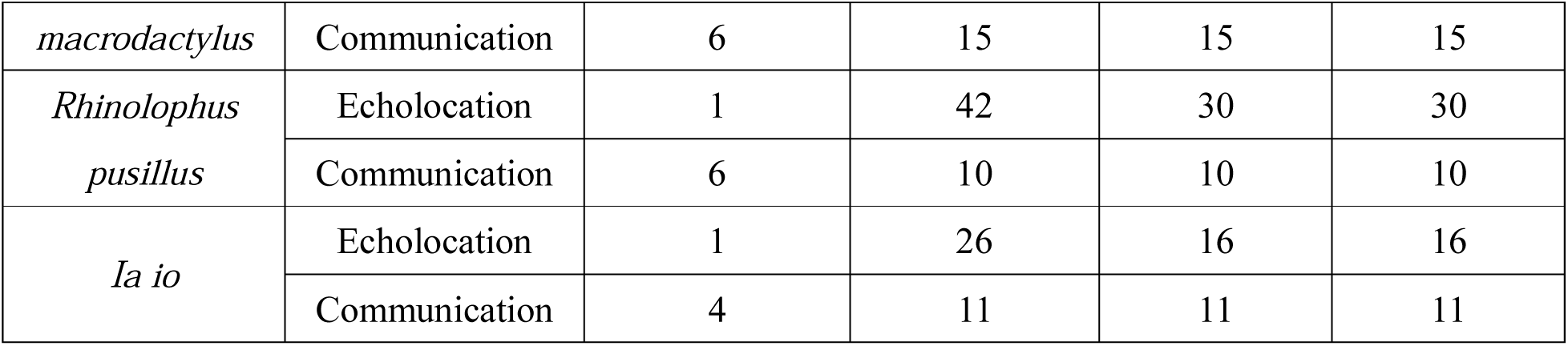
Separation results.

**Fig 1.**
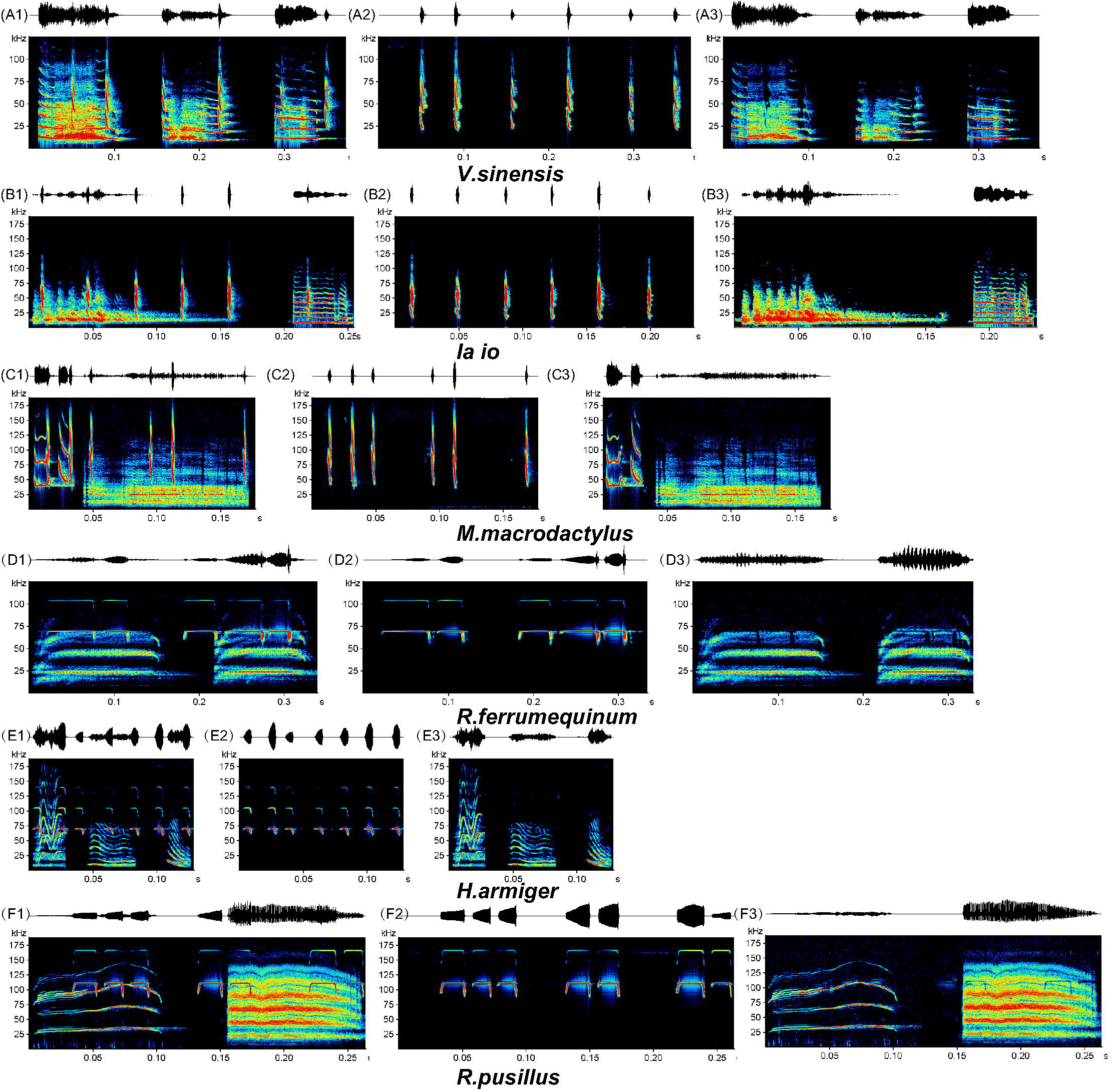
Spectrograms from original recordings of overlapping calls and calls separated by the BLSTM network. The first graph represents each line of the original overlapping calls and the second and third graphs show the separated echolocation and communication calls, respectively.

A comparison of seven temporal-spectrum parameters from the separated calls and the original recorded non-overlapping calls showed no significant differences (Fig 2 and S3 Table). In addition, parameter deviations in separated calls and original non-overlapping calls showed minimal RMSE values for both echolocation and communication signals (Fig 3 and Fig 4). Clustering analysis performed with separated echolocation calls produced an accuracy of 93.8% across species (Fig 5).

**Fig 2.**
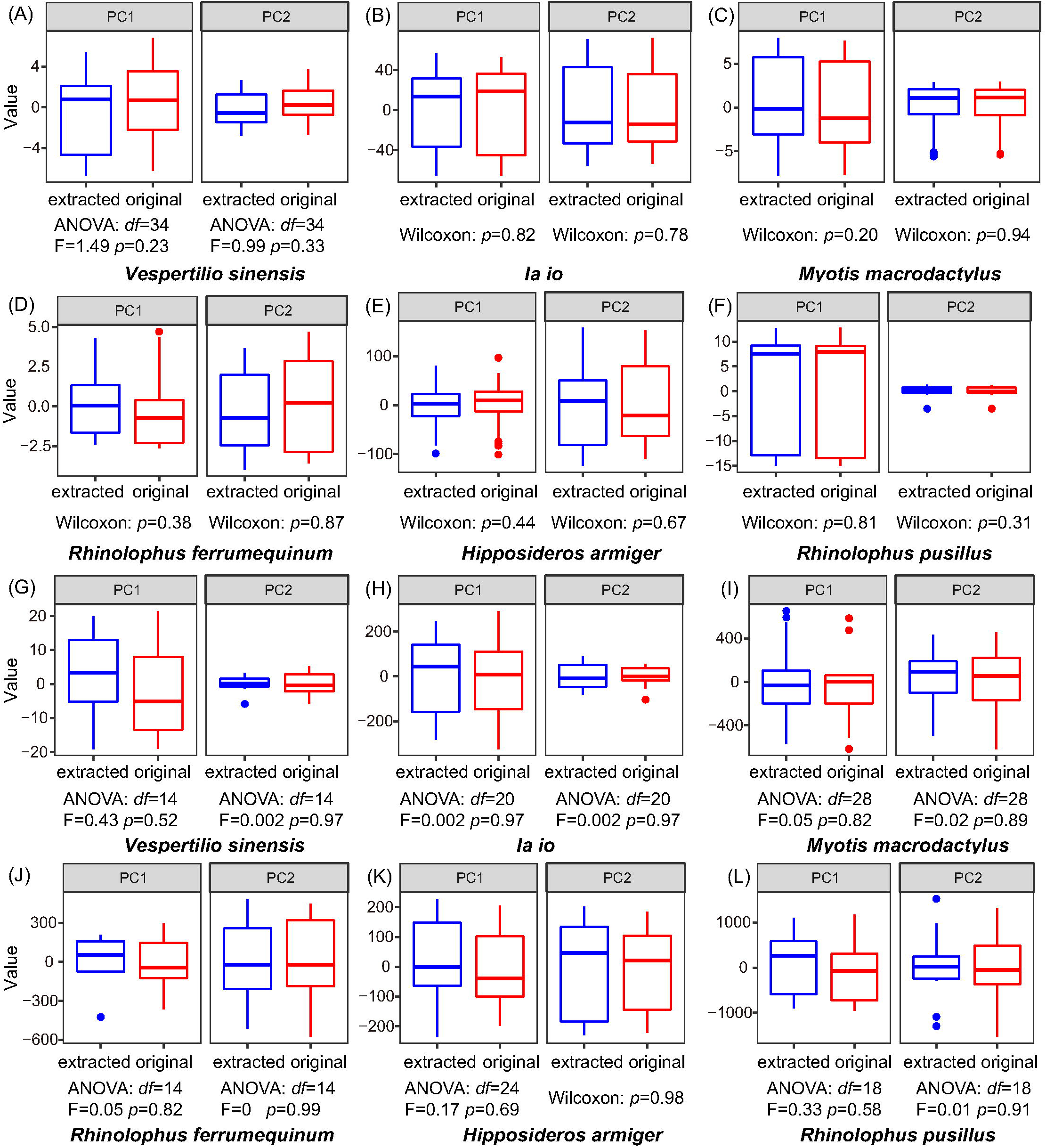
Comparisons between the separated and original calls. Two principle components extracted from seven temporal-spectral parameters were used in the study. Results for echolocation and communication calls are shown in (A-F) and (G-L), respectively.

**Fig 3.**
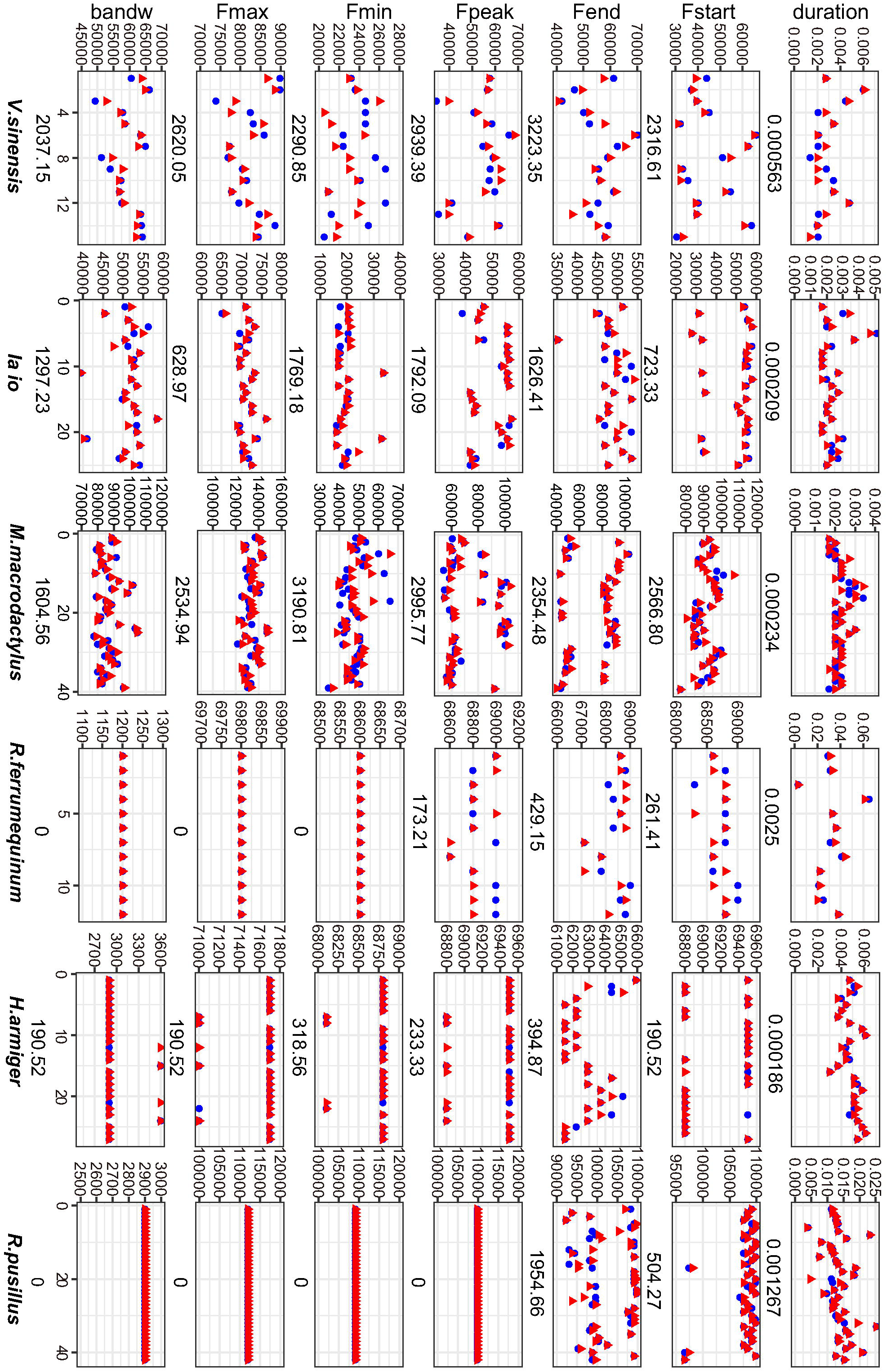
A comparison of deviations for separated and original echolocation calls. The RMSE value is shown under each plot. The vertical axis represents values for each parameter and the horizontal axis represents the number of syllables measured. The red triangles represent separated calls and the blue dots represent original calls. Abbreviations include duration (duration), Fstart (starting frequency), Fend (ending frequency), Fpeak (peak frequency), Fmin (minimum frequency), Fmax (maximum frequency), and bandw (bandwidth).

**Fig 4.**
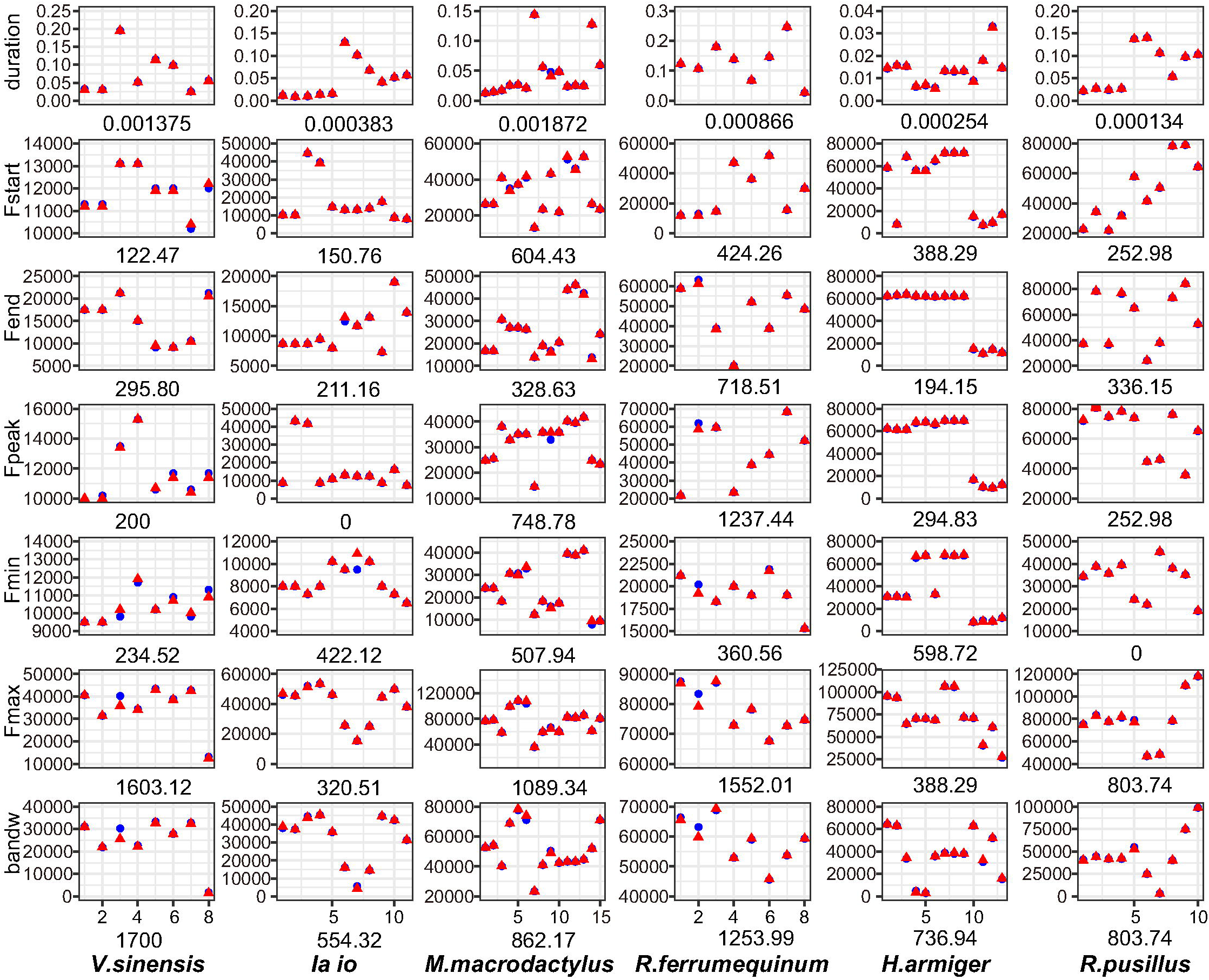
A comparison of deviations in separated and original communication calls. The RMSE value is shown under each plot. The vertical axes and abbreviations are the same in Fig 3.

**Fig 5.**
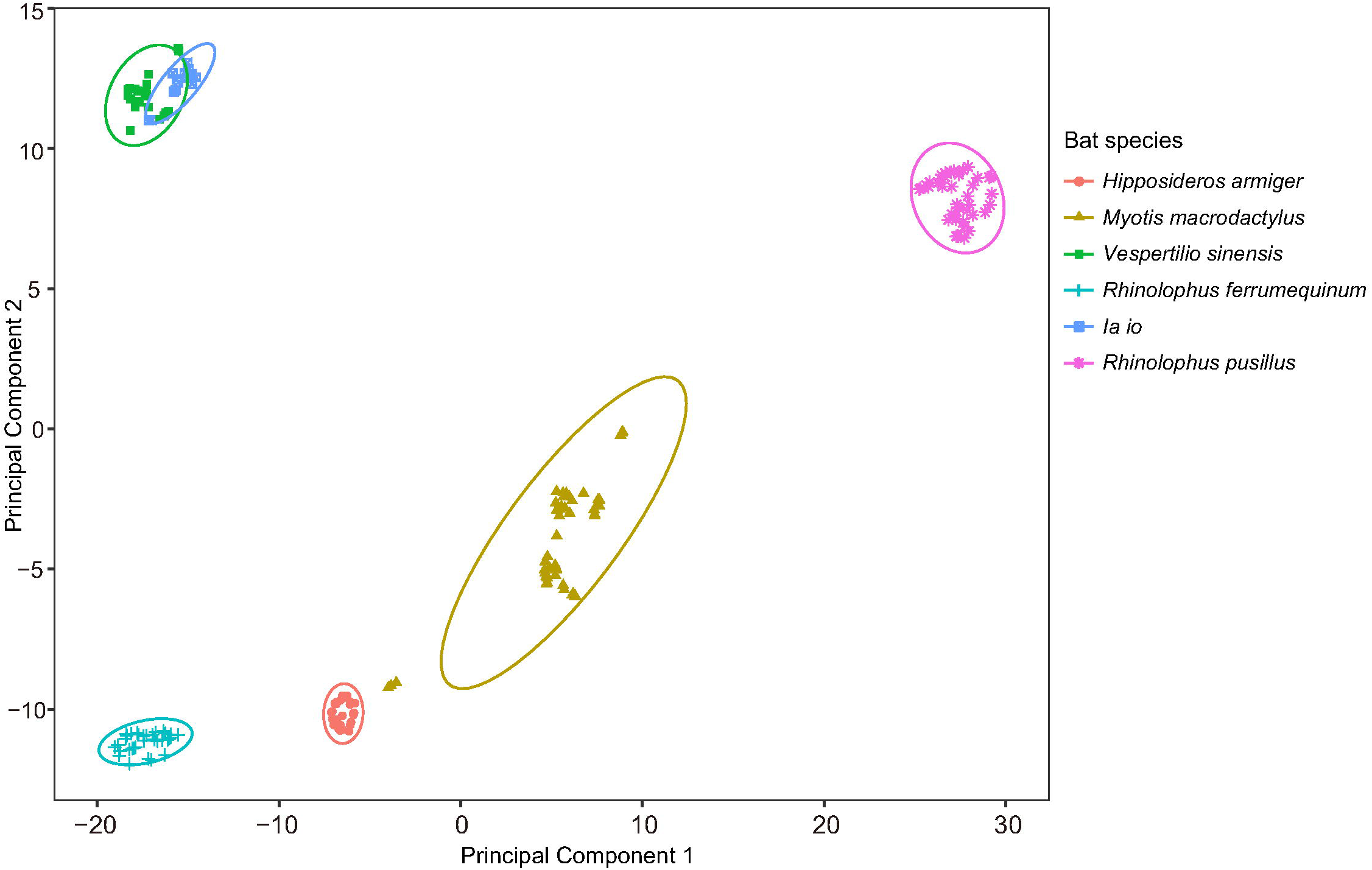
Clustering analysis for six bat species based on their separated echolocation calls. Overlapping echolocation signals cannot be used for species identification until after separation.

## Discussion

The BLSTM network used in the present study achieved high accuracy in separating overlapping echolocation and communication calls from bats. The training and validation loss for the model also exhibited fast convergence and high robustness for bat vocalizations. In particular, the separated calls extracted by the proposed algorithm were reconstructed as waveform files with nearly the same quality as the non-overlapping calls, suggesting BLSTM networks to be useful tools for separating signals in future bioacoustic research, such as sound analysis, acoustic identification, species classification, and wild animal monitoring.

It was difficult to compare the performance of this algorithm with that of previous studies, primarily because of differences in the experimental procedure. However, a comparison of temporal-spectrum parameters between separated calls and non-overlapping calls was included as an evaluation metric. The seven parameters used in this study are commonly used in bat studies to describe the temporal-spectral features of syllables [26, 29]. Statistical results for this comparison showed no significant differences and small deviations in parameters between separated calls and original recordings, indicating the system was able to separate calls without affecting syllable quality. In addition, clustering analysis conducted with reconstructed echolocation calls was highly accurate (93.8%) for species classification, indicating that calls separated from overlapping signals could be used to synthesize initial data.

The BLSTM network exhibited good performance across all six bat species using both narrow and broad time-frequency calls. It also successfully separated different syllable types from both overlapping echolocation and communication calls (Table 1, Fig 1). No species-specific *a priori* knowledge or particular acoustic sensor was directly encoded into the system, making it generalizable to other animal populations with additional training data. Although the dichotomy between communication and echolocation calls is relatively drastic, the proposed separation system has potential applications for other species, as such mixtures are very common in bats. In the future, more complex emitter-independent separation could be conducted using the proposed system, such as combinations of echolocation or social calls from other animals.

While deep learning models generally perform better when provided with more data, training with bat calls requires fewer samples than human speech separation, in which available training sets can exceed hundreds of hours [13]. One possible reason for this may be the high signal-to-noise ratio (SNR) of bat sounds recorded with high-quality ultrasound devices. Previous studies have indicated that a high SNR can improve separation accuracy [30] and our results suggest this model was suitable for use with small, high-quality datasets. Although the sound data in this study were sampled in controlled lab conditions, producing recordings that were essentially free of background noise, acoustic analysis software could potentially optimize the separation further by excluding any background noise that was present in the signal.

Future studies will also assess the performance of this network for other animal species. Stereotypical patterns and clearly classifiable syllables have been observed in the vocalizations of birds, non-human primates, whales, dolphins, and several other species [31-33]. Features used in the proposed BLSTM were log spectral magnitudes, which can be acquired from any vocal sound. This could potentially lead to robust software that is not specific to a certain species or task. The model could also be generalized to other animals, though limitations may exist. In addition to the quality and quantity of training samples, hyper-parameters must be tuned in accordance with the data [34, 35].

## Conclusion

A sound separation model was proposed for extracting bat calls, achieving excellent results. This is the first experimental evidence that the BLSTM model is suitable for separating overlapping bioacoustic signals. These results provided a new source for sound data analysis in animal acoustics research, which may contribute to sample sizes and improve efficiency. This study also demonstrates the potential of deep neural networks for applications to animal vocalization research, including species classification and speech separation.

## Materials and Methods

### Sound recording and data preparation

#### Species selection and sound sources

Echolocation calls from bats are primarily composed of constant frequency (CF) components and frequency modulated (FM) components. Social calls are composed of CF, FM, and noise-burst (NB) components. FM calls have short pulse durations and wide bandwidths. As such, they overlap with social calls less in time but more in frequency. In contrast, CF calls have long pulse durations and narrow bandwidths. They overlap with social calls more in time but less in frequency. In consideration of the varied overlapping patterns found in bat calls, we selected both CF bats (*Rhinolophus ferrumequinum, Hipposideros armiger*, and *Rhinolophus pusillus*) and FM bats (*Vespertilio sinensis, Myotis macrodactylus*, and *Ia io*) to test the separation capabilities of the proposed network, including six different species to test method generalizability.

Source sound files from *V. sinensis, M. macrodactyllus, R. ferrumequinum, R. pusillus*, and *H. armiger* were collected from previous studies in our lab (S1 Table). Sound files for *Ia io* were selected from unpublished data as follows. Bats captured from the field were housed in a husbandry room with abundant food and fresh water. During each sound recording experiment, 4–5 bats were transferred to a temporary cage. Sound recordings were collected using the Avisoft UltraSoundGate 116H (Avisoft Bioacoustics, Berlin, Germany) and a condenser ultrasound microphone (CM16/CMPA, Avisoft Bioacoustics). The sampling frequency was set to 375 kHz at 16 bits. The recording experiment lasted five days in order to acquire a sufficient number of recordings, beginning at 18:00 and finishing at 6:00 the following morning. S1 Table shows sample numbers and locations for the bats, as well as the total duration of sound files selected for the study. All experimental procedures complied with the ABS/ASAB guidelines for the Use of Animals in Research and were approved by the Committee on the Use and Care of Animals at the Northeast Normal University (approval number: NENU-W-2010–101).

#### Sound analysis

The total duration of recorded sound files (i.e., original recording files) used for each bat species is shown in S1 Table. We employed Avisoft-SASLab Pro (Version 5.2.12, Avisoft Bioacoustics, Berlin, Germany) to identify non-overlapping and overlapping syllables in echolocation and communication calls. These syllables and calls were described and classified following the nomenclature developed by Kanwal, Matsumura (36) and Ma, Kobayasi (37). The recorded non-overlapping calls were used for preparing training files of each call type and the recorded overlapping calls were used for separation.

#### Data preparation

Supervised machine learning algorithms use training samples to “learn” the steps required for completing a task. The training phase in this study involved preparing clear and non-overlapping echolocation and communication calls, selected from original recording sounds. In this process, the BLSTM network learned features found in both call types.

Training samples consisted of randomly selected non-overlapping syllables in echolocation and communication calls from each bat species (in the original recordings), with signal-to-noise ratios (SNRs) above -20 dB. The echolocation training files contained 1,300–6,240 pulses and the communication training files contained 780–1,800 syllables (S1 Table). Although the quantity of selected syllables varied between studies, the data was sufficient for model training. Efforts were made to include roughly equivalent quantities of each syllable type. Time intervals between syllables in the training files were consistent with those of the original recordings. The lengths of training files for echolocation and communication calls were the same for each bat species (S1 Table).

### Model training and call separation

#### Model structure and training stage

We developed a network with four BLSTM layers, followed by one feedforward layer (Fig 6). Each BLSTM layer included one forward and one backward basic LSTM layer, both of which were added with dropout functions (tensorflow.nn.rnn_cell.DropoutWrapper). Each BLSTM layer contained 300 hidden cells and the feedforward layer corresponded to the embedding dimension (i.e., a 3D matrix with depth N=40 in this experiment). Stochastic gradient descent with a momentum of 0.9 and a fixed learning rate of 10^−3^ was used for training. The tanh activation function and the Adam optimizer were adopted to support adaptive learning rates and faster convergence. The structure and hyper-parameters for the model were designed based on the work of Hershey, Chen (21).

**Fig 6.**
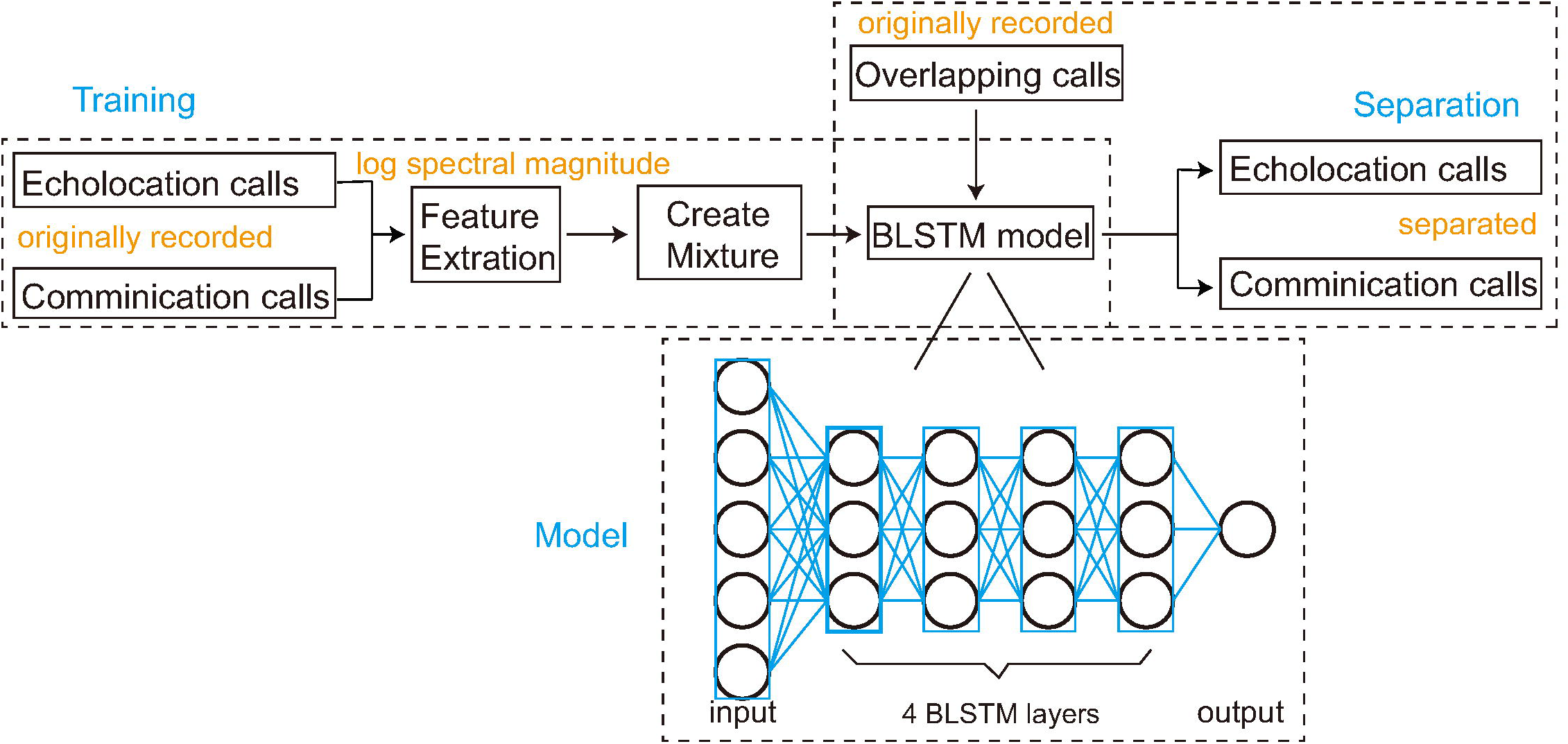
The BLSTM model architecture and workflow graph.

The model was trained using the files for one bat species in each trial. Echolocation and communication call training files were loaded using the librosa (version 0.6.2) Python package. Frames from the two sound files were read and added together to create sound mixtures. Sound features used for training (log spectral magnitudes) were extracted from this mixture. The extraction process was completed using a short-time Fourier transform (STFT) with a Hamming window (length of 512 and shift of 256).

The mixture from each bat species was then segmented into 100-frame samples, all of which were divided into a training set and a validation set using a ratio of 2:1 (see S1 Table for detailed sample quantities). The training set, validation set, and indicator labels were combined and input to the model. The validation set was used to optimize tuning parameters and evaluate call separation performance. Indicator labels were set to 0 or 1, representing the two types of calls in the mixture. Ideal binary masks were used to train the network and gradients were calculated using shuffled mini-batches (batch size of 128) from larger segments.

The output of this model was a set of embeddings that included learned features for both echolocation and communication calls. In this framework, the deep network assigned embedding vectors to each time-frequency bin in the spectrogram. The network then minimized the distance between embeddings dominated by the same call type in each bin while maximizing the distance between embeddings dominated by different call types. The output was then compared with the validation set and indicator labels to calculate loss, which was back propagated from the output to the input through each layer. Model weights and parameters were then updated based on the calculated loss and training was completed after sufficient iteration epochs.

#### Separation stage

In this stage, overlapping echolocation and communication calls were randomly selected from the original recordings to create a sound file of test sets, used for separation. The log spectral magnitudes of the overlapping calls were then extracted, combined into samples, and input to the trained model. The phases of calls extracted from the sound files were also saved for use in sound reconstruction. The trained model then output embeddings for each segment (100 frames) in a process similar to the training stage. Embeddings were clustered using the k-means method from Scikit-learn (Version 0.20.0) to produce time-frequency masks. The number of clusters corresponded to the number of call types in the mixture (2 - echolocation and communication). These masks and the clustering method were then used to determine which parts of each segment in the overlapped calls would be preserved or neglected based on their correspondence to each call type. For example, if the maximum magnitudes were more likely to belong to echolocation calls, the related mask values were set to 1 and the others were set to 0, allowing the echolocation calls to be separated correctly. Finally, output calls were reconstructed using the inverse fast Fourier transform (IFFT) function numpy.fft.ifft in NumPy (Version 1.15.1). The IFFT transformed the magnitude into a wave using phase information saved at the beginning of the separation stage. The model produced two waveform files, each containing one call type. Additional detail concerning the sound separation algorithms can be found in the work of Hershey (2016).

### Model evaluation

The quality of reconstructed echolocation and communication calls was assessed by comparing their temporal-spectrum parameters to the non-overlapping calls selected from the original recording files (excluding training data). Avisoft-SASLab Pro was used for automatic parameter measurements of duration, bandwidth, peak frequency, minimum frequency, maximum frequency, starting frequency, and ending frequency. A t-SNE (t-distributed stochastic neighbor embedding - R3.6.1 package) analysis was adopted for dimensionality reduction. Two dimensions were extracted from these seven parameters for original and separated syllables and compared with one-way ANOVA (aov in R3.6.1) or two-sided Wilcoxon signed-rank tests (wilcox.test in R3.6.1), depending on their fit to a normal Gaussian distribution. The significance level was set to 0.05 for all tests. We adopted the root mean square error (RMSE) to measure and avoid obscuring individual variations between reconstructed and original calls. Clustering analysis was conducted using the reconstructed echolocation calls from the six bat species, to assess whether the separated calls could be further used in species classification.

## Acknowledgements

We are grateful to Dr. Yanhong Xiao of the Experimental Center of the School of Environment at Northeast Normal University, for her assistance in acquiring the experimental materials. We thank LetPub (www.letpub.com) for its linguistic assistance during the preparation of this manuscript.

## Author contributions

**Conceptualization**: Kangkang Zhang, Walter Metzner.

**Data curation**: Tong Liu, Shengjing Song, Xin Zhao.

**Formal Analysis**: Kangkang Zhang, Tong Liu.

**Methodology**: Kangkang Zhang, Ying Liu, Jiang Feng.

**Software**: Kangkang Zhang, Shijun Sun.

**Funding acquisition**: Ying Liu, Jiang Feng, Walter Metzner.

**Supervision**: Jiang Feng.

**Visualization**: Kangkang Zhang, Tong Liu.

**Writing – original draft**: Kangkang Zhang, Ying Liu.

**Writing – review & editing**: Ying Liu, Walter Metzner.

## Supporting information

**S1 Table. A summary of calls used for model training.**

**S2 Fig.**
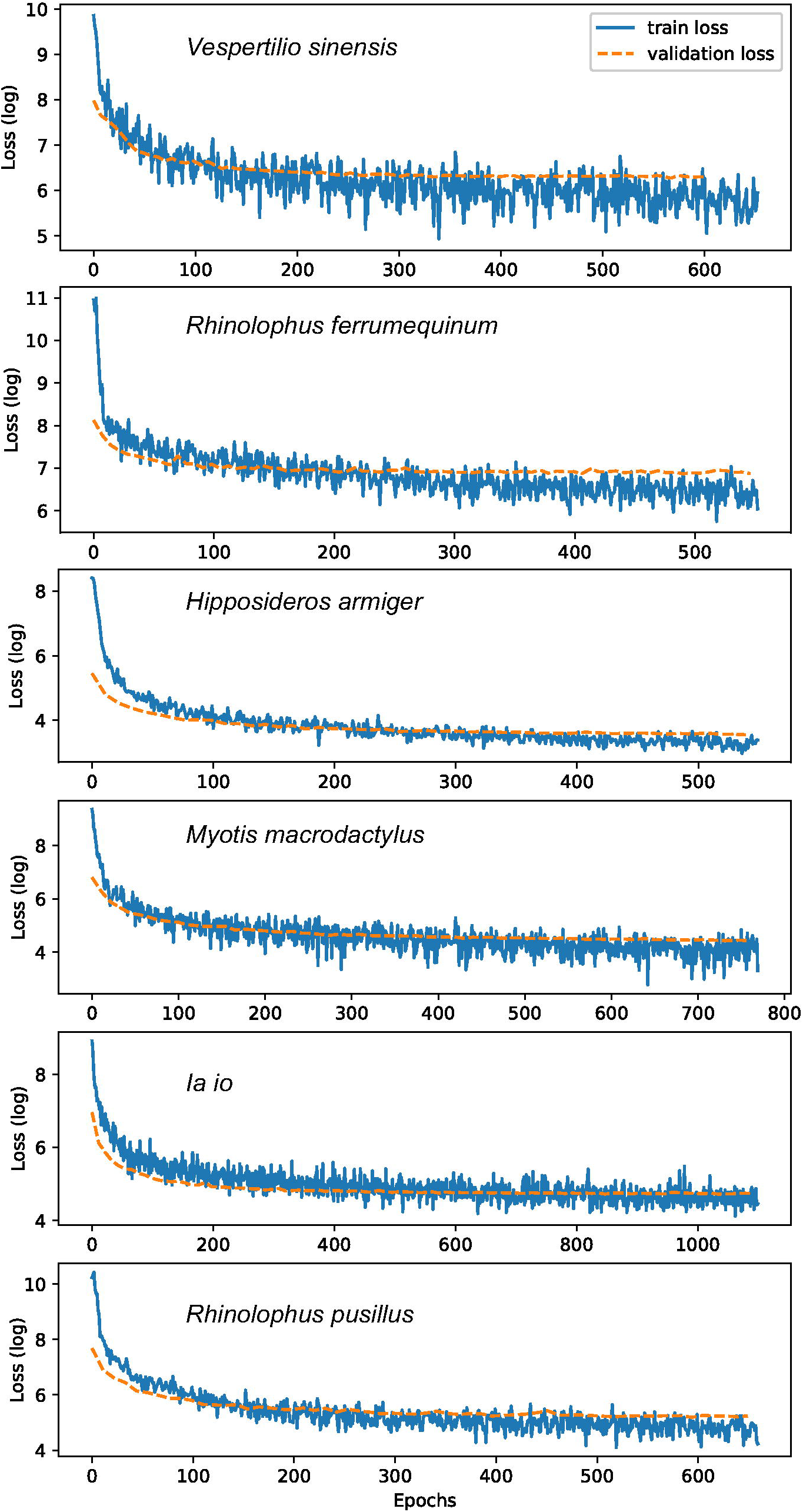
Training loss and validation loss during model training.

**S3 Table. Statistical comparisons of principle components extracted from seven parameters.** No significant differences were observed between parameters for separated and original syllables. A one-way ANOVA was used to test the normal distributed data and a two-sided Wilcoxon signed-rank test was used to assess the data that did not conform well to a normal distribution.

