## Supplemental Material S3 Table for "Separating overlapping bat calls with a bi-directional long short-term memory network"

| Types of calls | Species | Parameters | Statistics | df | F | P |
| --- | --- | --- | --- | --- | --- | --- |
| Echolocation | *V.sinensis* | PC1 | ANOVA | 34 | 1.494 | 0.23 |
|  |  | PC2 | ANOVA | 34 | 0.989 | 0.327 |
|  | *Ia io* | PC1 | Wilcoxon | 48 |  | 0.823 |
|  |  | PC2 | Wilcoxon | 48 |  | 0.778 |
|  | *M.macrodactylus* | PC1 | Wilcoxon | 106 |  | 0.203 |
|  |  | PC2 | Wilcoxon | 106 |  | 0.936 |
|  | *R.ferrumequinum* | PC1 | Wilcoxon | 22 |  | 0.384 |
|  |  | PC2 | Wilcoxon | 22 |  | 0.87 |
|  | *H.armiger* | PC1 | Wilcoxon | 52 |  | 0.441 |
|  |  | PC2 | Wilcoxon | 52 |  | 0.672 |
|  | *R.pusillus* | PC1 | Wilcoxon | 82 |  | 0.809 |
|  |  | PC2 | Wilcoxon | 82 |  | 0.308 |
| Communication | *V.sinensis* | PC1 | ANOVA | 14 | 0.426 | 0.524 |
|  |  | PC2 | ANOVA | 14 | 0.002 | 0.966 |
|  | *Ia io* | PC1 | ANOVA | 20 | 0.002 | 0.969 |
|  |  | PC2 | ANOVA | 20 | 0.002 | 0.973 |
|  | *M.macrodactylus* | PC1 | ANOVA | 28 | 0.052 | 0.821 |
|  |  | PC2 | ANOVA | 28 | 0.019 | 0.891 |
|  | *R.ferrumequinum* | PC1 | ANOVA | 14 | 0.051 | 0.824 |
|  |  | PC2 | ANOVA | 14 | 0 | 0.993 |
|  | *H.armiger* | PC1 | ANOVA | 24 | 0.169 | 0.685 |
|  |  | PC2 | Wilcoxon | 24 |  | 0.98 |
|  | *R.pusillus* | PC1 | ANOVA | 18 | 0.325 | 0.576 |
|  |  | PC2 | ANOVA | 18 | 0.012 | 0.914 |

**S1 Table. Statistical comparisons of principle components extracted from seven parameters.**

No significant differences were observed between parameters for separated and original syllables. A one-way ANOVA was used to test the normal distributed data and a two-sided Wilcoxon signed-rank test was used to assess the data that did not conform well to a normal distribution.
